# An Affordable and Customizable Arduino-based Thermo-controller for Thermal Ecology Experiments

**DOI:** 10.1101/2024.08.23.609058

**Authors:** Cassidy Hawk, Erik Iverson, Justin Havird

## Abstract

Insight into organismal physiology, evolutionary ecology, and climate change susceptibility all rely on high-quality and easily-obtainable thermal tolerance information. A variety of methods exist for measuring thermal tolerance, and thermal tolerance metrics are highly influenced by small differences in methodology. To promote consistency, reproducibility, and accessibility, we present here instructions for building an Arduino-based thermo-controller for raising, lowering, and holding temperature in a variety of experimental conditions. The setup is affordable (∼$100), highly customizable, and highly consistent in the outcomes it produces. We include data from real thermal trials demonstrating the ability of the system to meet, hold, and reproduce desired ramping rates and hold temperatures.

## Introduction

Studies of thermal limits and performance at various temperatures are essential to our understanding of basic organismal biology as well as to applications such as predicting species’ responses to climate change. Among thermal ecologists, the most foundational metrics are the critical thermal maximum (CT_max_) and minimum (CT_min_) of ectotherms, which define the tolerable temperature range on short time-scales, and the upper and lower critical temperature (UCT & LCT) of endotherms, which define the thermoneutralzone (TNZ) over which metabolic rate is constant. These parameters have been measured for hundreds of species, permitting large-scale phylogenetic assessments of how these traits evolve (Bennett et al., 2021; Comte & Olden, 2017; N. L. Payne et al., 2021). Additionally, within species, these traits might be measured for hundreds of individuals, allowing for the identification of the genetic variants responsible for thermal tolerance (C. Payne et al., 2022). Thermal traits like these are determined not just by genetics, but by the early life environment and the acclimation conditions of the organism, and measurement of critical limits after acclimation to various temperatures is a common dimension of study (Ørsted et al., 2022). Untangling the genetic and environmental controls on thermal tolerance is a key aspect of understanding range limits and interspecific competition, and in predicting how and why species will be affected by rising temperatures.

While critical thermal measurements are one of the most widely used traits in ecology, they are notoriously sensitive to experimental conditions, making repeatability and consistency key (Desforges et al., 2023; Leong et al., 2022; Ørsted et al., 2022). For instance, robust and reliable estimates of thermal performance parameters require, among other things, consistent, smooth temperature ramping and/or accurate and reliable temperature holding. While a variety of schemes have been used to achieve desired ramp rates or hold temperatures, including even gas stoves (Culumber et al., 2012), the ideal apparatus involves an automated thermo-controller which compares the realized temperature or rate to the desired state and adjusts temperature accordingly (i.e., a thermostat). Given the wide variety of apparatus that may be used, the unique needs of each organism, and the many different methods for raising or lowering temperature, an ideal thermo-controller would be adaptable to many different experimental situations. Such a device would also be simple and cheap, given the need to drastically expand experimental efforts in the world’s most biodiverse countries, where dedicated scientific funding and equipment is lacking. Here, we present a thermo-controller than can be built and operated by anyone without specialized electrical or programming knowledge for less than $100 with parts readily available online. This customizable controller can ramp temperature both up or down by gating an electrical circuit to which a variety of heaters, pumps, or chillers can be connected, and can also hold temperature constant. The ramping rate and hold temperature are set by the user, and the device also has a button for logging when individual animals in a trial reach their CT_max_ or CT_min_, or noting any other behavior the user desires in an electronic format. We provide temperature data from a variety of experiments demonstrating how the controller performs.

## Materials and Methods

### Overview of construction and programming

The thermo-controller is based around an Arduino Uno board (**Supplement S1** Supplies List) which allows easy construction of integrated circuits without soldering (**Supplement S2** Wiring Guide). Housed inside a water-proof box, the board controls the flow of power from a wall socket to a 3-outlet power strip to which heaters, pumps, chillers, or any other temperature-changing apparatus can be connected. A light on the waterproof housing signals to the user whether power is flowing or interrupted. The board has two inputs, a thermometer to assess realized temperature and facultatively turn the flow of power on or off, and a button to record the time and temperature at which animals reach their CT_max_ or CT_min_. Data is logged to a PC via a USB cable and can be displayed in real time. We provide three starter scripts (**Supplement S3** Quick Start Guide) for CT_max_, CT_min_, and a constant-temperature hold.

In a CT_max_ protocol, the user determines the desired ramping rate in degrees Celsius / minute (i.e., 0.3 °C / min) and the length of the window (in seconds) to use for calculating the ramping rate. The controller samples temperature once per second and compares the average rate of temperature increase in the window (i.e., prior 40 seconds) to the desired ramping rate, and turns the power on if the real ramping rate is lower than the desired rate. If connected to a heating element, this facultatively turns the power of the heater on and off to match a desired rate. To begin, the user uploads code to the controller using Arduino IDE and then opens ArduSpreadsheet to view the time and temperature data in real time. When the controller has reached the set number of observations in the sliding window (i.e., 40 seconds), it will begin gating the power supply. The user then plugs the controller into external power, initiating temperature changes in the experimental chamber. When animals reach their CT_max_, the user presses the button on the box, creating a numbered entry in the spreadsheet displaying user-determined text as well as the current time and temperature. The light on the box helps the user determine if the system is functioning and raising temperature correctly. If the light alternates between on and off, then the controller is adjusting temperature as expected; if the light remains on constantly, the heating element is not adequate for raising the temperature quickly enough to meet the target rate, causing power to be constantly on. If the light is off for a period, the system has overshot the target rate and may be heating the organism faster than desired. The real-time logging of temperatures in ArduSpreadsheet allows the user to adjust or abort the protocol before animals reach their CT_max_ if the ramping rate is not correct. A CT_min_ protocol works similarly except that the user runs a different script, sets a negative ramping rate, and connects a cooling element such as a chiller, refrigerator, or a pump connected to an ice water bath. A temperature-hold protocol turns power on if the current real temperature is below or above the desired temperature.

### CT_max_ setup

The controller can be used with any heating element to perform CT_max_. The actual ramping rates achieved will differ from the desired rate somewhat based on the type of heating element used and the volume and medium that are being heated. We tested its performance with a double water bath setup and aquarium heaters for aquatic organisms, which we describe here. Two 800w Aquarium Heaters (TH-0800S Deluce Titanium tube, Finnex, Chicago, Illinois, USA) were connected to the controller’s outlets, and two 80GPH circulation pumps (Uniclife 80 GPH Submersible Water Pump) were connected to constant power and placed in a ∼10L open top external water bath (Figure 2A). An internal tank was placed in the center and filled with preconditioned water (matching organismal acclimation conditions) as well as two air stones to keep the water oxygenated. Water in the outer chamber was filled to below the inner chamber, and can be from any source as organisms do not come into contact with it. Water in both the inner and outer chambers should be at the organism’s acclimation temperature. The controller’s thermometer was placed into the center of the internal tank, affixed so that it was at least 5 cm from any wall, and the 800w heaters were plugged into the controller’s outlets. CT_max_ temperature profiles were analyzed to calculate real ramping rates (Results).

### CT_min_ setup

Running CT_min_ with the thermo-controller is similar to CT_max_ except that a different script is used and a cooling element is connected rather than a heating element. In the case of a water bath setup for aquatic organisms, this might mean controlling a pump that either passes water from the external tank through an electric chiller or refrigerator, passes it through a heat-transfer coil submerged in an ice bath, or directly exchanges water between an ice bath and the external chamber. We tested performance using the latter two methods, with a 400 GPH pump plugged into the controller used either to sending water out of the external tank, through a coiled metal heat sync, and back into the external tank, or to pump water directly from the ice bath into the external chamber and return water to the ice bath via overflow. Two 80GPH circulation pumps connected to constant power were used as in CT_max_ to move water around in the external bath.

As the pump exchanges heat between the external chamber and the ice bath, the temperature differential between them decreases; this makes it hard to continue to achieve the desired cooling rate at low temperatures (< 10 C) under either the heating coil or direct water transfer method.

This causes the controller to remain on constantly while achieving little. To overcome this, we added ice packs (frozen 50 mL falcon tubes) directly to the inner chamber at low temperatures, using the light on the controller as a guide to whether the intervention was lowering temperatures quickly enough.

### Temperature-hold setup

We performed a temperature-hold protocol with a water-bath using the same setup and heaters as in CT_max_ to keep water at a constant temperature above ambient. We did not extensively trial the protocol for holding water to a cooler temperature than ambient; however, this should run similarly to the other hold protocol but with the cooling elements from CT_min_ being used instead of heaters. One caveat is that adding cold water from an ice bath, which needs time to circulate, is likely to result in cooling overshoots, and it may take a very long time for the temperature to rise to ambient again. This is likely to result in a much more variable temperature than when electric heaters are used to raise the temperature above ambient.

## Results

Here we report the results of 17 CT_max_ runs, 17 CT_min_ runs, and 17 above-ambient temperature hold runs using the setups described above. For CT_max_ and CT_min_, the realized temperature ramping rate is not directly analogous to the user-set temperature ramping rate. Rather, it is a function of the volume of water used in both chambers, the surface area of contact between the chambers, the thermal conductivity of the inner chamber wall, the number of heating or cooling elements, the power of heating or cooling elements, the responsiveness of heating or cooling elements to being turned on, and the number of temperature readings being averaged, and the ambient temperature in the room. Thus, for the particular setup of any user, some trial-and-error is required to determine what combination of set temperature and averaging will produce the desired heating rate.

### CTmax runs

We determined that with the setup described above, temperature averaging over 40 seconds and a set ramping rate of 0.6 approximately resulted in the desired ramping rate close to 0.3 °C / minute, a fairly standard ramping rate in the eco-physiology literature. Over 17 runs in these conditions, the average ramping rate in the rising leg of the trial was 0.332 ± 0.003 SE °C / minute. This steady ramping rate was achieved after roughly 10 minutes during which the rate increased from 0 to the target rate. Temperature rose smoothly and steadily to temperatures in excess of common aquatic organismal thermal tolerance limits (> 40 °C).

### CT_min_ runs

We used the setup described above, temperature averaging over 40 seconds, and a set ramping rate of -0.4 °C / minute over 17 runs to achieve an average ramping rate in the falling leg of a CT_min_ trial of -0.238 ± 0.009 SE °C / minute. This steady cooling leg began more quickly, after approximately 5 minutes or less, due to the mechanism of injecting already cooled water into the outer bath. This also caused ramping to be less smooth overall. Large pulses were frequently followed by periods of inactivity, producing a jerkier, staircase graph of temperature change. Below ∼12 °C, ramping began to lag the desired rate as temperatures became more equivalent between the external chamber and the ice bath, necessitating the use of ice packs in the internal chamber.

### Temperature-hold runs

Using the setup described above, the controller accurately matched the exact hold temperature and no modification of the setting number was needed. Trials ranged from 20 minutes to over an hour. The average temperature over all trials at 1-second intervals with a set temp of 18 °C was 18.138 ± 0.068 SE °C, with an average within-trial standard deviation of 0.103 ± 0.025 SE °C (n = 11 trials). When set to 24 °C, the average temperature was 23.937 ± 0.023 SE °C, with an average within-trial standard deviation of 0.041 ± 0.005 SE °C (n = 11). When set to 30 °C the average temperature was 29.943 ± 0.035 SE °C, with a within-trial standard deviation of 0.120 ± 0.030 SE °C (17 trials).

## Discussion

We demonstrated that the thermo-controller described here, which can be built without specialized electronics or programming knowledge for ∼$100 with readily available parts, can perform CT_max_, CT_min_, and temperature-holding procedures with a high degree of customizability and precision. This makes the device accessible to researchers with limited resources for a variety of applications in eco-physiology. Unlike fixed-input heating or cooling methods such as burners or refrigerators, this thermo-controller is easily adjustable to specific of heating or cooling rates and accommodates setups with varying levels of water volume, conductivity, and power. The system also possesses several features useful to experimenters, including the ability to display temperature data in real time, the ability to assess the power status of the device and adjust experimental conditions if necessary, and the ability to log events such as CT_max_ and CT_min_ endpoints with a button. In our trials, temperature ramping rates could be achieved with a high degree of precision, deviating little (average SE) between trials or over the duration of the trial. The temperature hold protocol was even more precise, deviating only by fractions of a degree over the course of trials. This makes the system described here an affordable, reliable, and useful experimental setup for the kinds of thermal tolerance and metabolic experiments needed in a variety of species to evaluate their susceptibility to climate change.

## Supporting information

Supplement S3 Quick Start Guide

Supplement S2 Wiring Guide

Supplement S1 Supply List

Data for runs analyzed

## Acknowledgments

We thank members of the Havird Lab at and Ryan Lab at UT Austin as well as the Schumer Lab at Stanford University for building and testing prototypes of this device.

