## Supplement S3 Quick Start Guide for "An Affordable and Customizable Arduino-based Thermo-controller for Thermal Ecology Experiments"

An Affordable, Customizable, and Consistent Arduino-based Thermocontroller for Measuring Critical Thermal Maximum & Minimum (CT_max_ & CT_min_)

Cassidy Hawk, Erik Iverson, and Justin Havird

Step 1: Download “Arduino IDE”, a software that can be found on the Arduino website.

Step 2: Download the code used to control the Arduino: <https://github.com/Hawkpun/Critical-Thermal-Tolerance-Code>

Step 3: Open the code downloaded using the Arduino IDE software, and using the Library Manager found on the left side of the window, install *OneWire* by Jim Studt and *DallasTemperature* by Miles Burton.

Step 4: Plug in the Arduino Uno to your computer and select it using Tools > Board >Arduino Uno. Additionally, navigate to Tools > Ports > select Arduino from the list.

Step 5: To manipulate the desired ramping rate, find and adjust “TempSet” in the code which by default is set to 0.3°C*min^−1^ .

Step 6: Upload the code to the Arduino and navigate to the Serial Monitor. A 40 second calibration timer will begin counting down before Rate (°C*min^−1^), Time (in seconds), and Temperature (in Celsius) are displayed.

Step 7: Data collection can be handled in a variety of ways, but if a PC is being used then *Microsoft Data Streamer* is the most user friendly. To set this up, ensure that the serial monitor in the Arduino IDE software is closed.

**Other Notes**

“TempSet” refers to the rate of temperature increase and may require additional calibration depending on environmental factors such as room temperature. This setup is also equipped with a button designed to aid in data collection. Pressing the button will record the following: “CTmax Reached”, Organism #, temperature, and time in seconds. Organism # will increase by +1 each time the button is pressed. “Organism #” can be customized to state whatever species is currently being tested. To do this, Ctrl+F to find *Serial.print("Organism #");*, and change the text within the quotation marks.
