## Supplement S2 Wiring Guide for "An Affordable and Customizable Arduino-based Thermo-controller for Thermal Ecology Experiments"

An Affordable, Customizable, and Consistent Arduino-based Thermocontroller for Measuring Critical Thermal Maximum & Minimum (CT_max_ & CT_min_)

Cassidy Hawk, Erik Iverson, and Justin Havird

**Construction**

The Arduino Uno consists of numerous ports that are clearly labeled on the device itself. It contains 5 analog jumper pin inputs (A1-A5), 14 digital ports (0-13), 3 ground ports (GND), and 3 dedicated power outputs (5V-3.3V). The construction of this device will rely heavily on this terminology and is shown in the schematic. The Arduino Uno in conjunction with the breadboard and jumper wires will result in a solder free building process.

Before beginning the wiring process, it is important to cut three appropriate holes in the project box and feed the thermometer, button, and extension cord through the cutouts.

Connect wires as shown in the wiring schematic. Red wires depict positive, black wires depict negative, and yellow wires are data ports. The pink stars indicate connection points. Every connection point not on the breadboard or the Arduino Uno are not compatible with jumper wires and will require stripping the wires and twisting them together. These can be secured with electrical tape or hot glue. Important note, the extension cord requires splicing to properly wire. There is a ribbed negative side and a smooth positive side. Cutting a pealing back the smooth positive side should reveal raw copper wire, which will then be connected to either side of the solid-state relay.

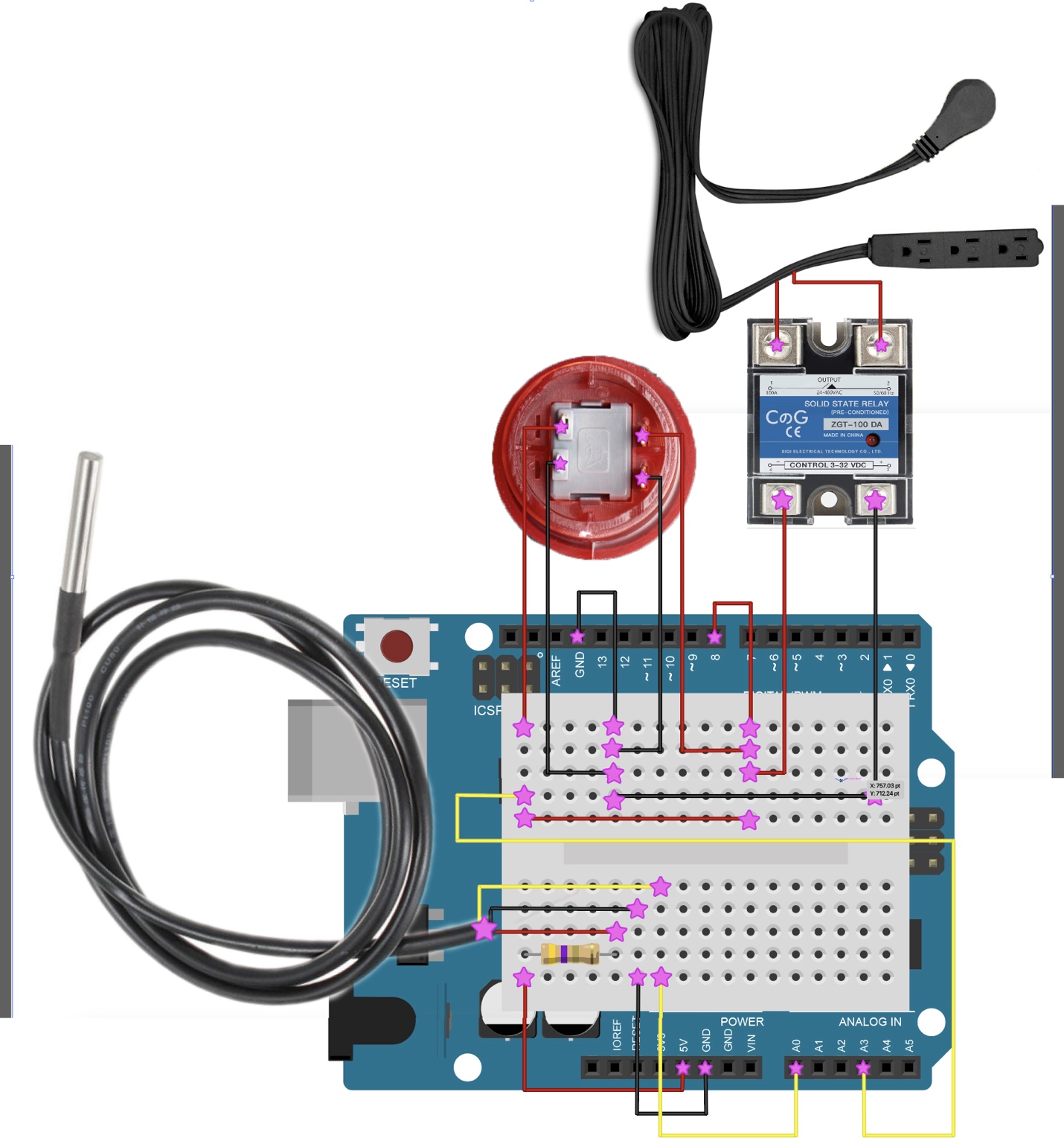
