## Supplement S1 Supply List for "An Affordable and Customizable Arduino-based Thermo-controller for Thermal Ecology Experiments"

**Materials**

Supplies List

Parts List:

- [Arduino Uno Rev3](https://store-usa.arduino.cc/products/arduino-uno-rev3/?gclid=CjwKCAjwwdWVBhA4EiwAjcYJEHIca7Pe1KuOvhQVY8Uat3Id9NCwojznly9rSFJ9F4D8fqPifyJ99BoCtmIQAvD_BwE)
- [Bread Board](https://www.adafruit.com/product/65)
- [(2x) Heating Element](https://www.amazon.com/Finnex-Deluxe-Titanium-Heating-Tube/dp/B0199W751Q/ref=asc_df_B0199W751Q/?tag=hyprod-20&linkCode=df0&hvadid=167149786275&hvpos=&hvnetw=g&hvrand=10510317476063400795&hvpone=&hvptwo=&hvqmt=&hvdev=c&hvdvcmdl=&hvlocint=&hvlocphy=9028280&hvtargid=pla-305107529430&th=1)
- [DS18B20 Temperature Sensor](https://www.adafruit.com/product/381?gclid=CjwKCAjwwdWVBhA4EiwAjcYJEK_Kl1H56p3UsjW766sWyP39WS0CguSvb1s3HsLLZCnl0pHu8aXqBhoCFegQAvD_BwE) (with 4.7k resistor)
- [USB 2.0 cable type a/b](https://www.adafruit.com/product/62?gclid=CjwKCAjwwdWVBhA4EiwAjcYJELoXRJ0crE1KN6FHZAw094lGkXO8Cew8Dx0Gk07Z3pGzWJqHM8jRSRoCPXIQAvD_BwE)
- [0.3 Meter USB Type B 2.0 Mount Cable](https://www.amazon.com/Kework-Extension-Motorcycle-Dashboard-Transferring/dp/B08P1CTJ65/ref=pd_ybh_a_36?_encoding=UTF8&refRID=XRN0SF8ZYT8RK3V12KSJ&th=1)
- [Male/Male Jumper Wires](https://www.adafruit.com/product/1956)
- [Button with LED](https://www.adafruit.com/product/3489)
- [200mm by 120mm by 75mm Waterpoof Project Box](https://www.amazon.com/Waterproof-Electronic-Plastic-Junction-Enclosure/dp/B06XSQZ5M6/ref=pd_ybh_a_21?_encoding=UTF8&psc=1&refRID=B9E5RH1D6ZDPPA2SYPPP)
- [Waterproof Joint Gaskets](https://www.amazon.com/dp/B01MDTUIIA/ref=sspa_dk_detail_1?pd_rd_i=B01MDTUIIA&pd_rd_w=TYHBI&content-id=amzn1.sym.3481f441-61ac-4028-9c1a-7f9ce8ec50c5&pf_rd_p=3481f441-61ac-4028-9c1a-7f9ce8ec50c5&pf_rd_r=3RJE14MTWAVMV7WX379P&pd_rd_wg=BIWIt&pd_rd_r=6d43c802-0c31-4a9f-b459-fe4dd470402d&s=hi&spLa=ZW5jcnlwdGVkUXVhbGlmaWVyPUFZWFZaNVNPNk5SR0cmZW5jcnlwdGVkSWQ9QTAxNTUyNzkyV1VNWEZTTTY3M0o3JmVuY3J5cHRlZEFkSWQ9QTA0NTY4MzkyT0lKQ00wR05CUldOJndpZGdldE5hbWU9c3BfZGV0YWlsX3RoZW1hdGljJmFjdGlvbj1jbGlja1JlZGlyZWN0JmRvTm90TG9nQ2xpY2s9dHJ1ZQ&th=1)
- [Solid State Relay (5V DC input / ~120V AC output)](https://www.amazon.com/SSR-100DA-3-32VDC-Output-24-480VAC-Plastic/dp/B08GP7Y292/ref=sr_1_2_sspa?keywords=solid%2Bstate%2Brelay&qid=1656086341&sr=8-2-spons&spLa=ZW5jcnlwdGVkUXVhbGlmaWVyPUEyQUZPRktHNzA2SUVFJmVuY3J5cHRlZElkPUEwOTI4NDgwM080RkxIQ01RNkxVWCZlbmNyeXB0ZWRBZElkPUEwNDMyOTg4MlFDWjJONEM1T0NFOSZ3aWRnZXROYW1lPXNwX2F0ZiZhY3Rpb249Y2xpY2tSZWRpcmVjdCZkb05vdExvZ0NsaWNrPXRydWU&th=1)
- [3 Prong Extension Cord with Multiple Outlets (Flat 3 ribbed cable)](https://www.amazon.com/BindMaster-Extension-Grounded-outlets-Angled/dp/B074P83P79/ref=sr_1_3_sspa?crid=3462M0KDE1KG8&keywords=3%2Bprong%2Bextension%2Bcord&qid=1656086659&s=industrial&sprefix=3%2Bprong%2Bextension%2Bcord%2Cindustrial%2C114&sr=1-3-spons&spLa=ZW5jcnlwdGVkUXVhbGlmaWVyPUEzSjZQR1IwUk5CWFM0JmVuY3J5cHRlZElkPUExMDQyNTk1MTlJVDNYNTlSRjFJTCZlbmNyeXB0ZWRBZElkPUEwMjQzNjQwMzlPVFUxOE1YOUo2TSZ3aWRnZXROYW1lPXNwX2F0ZiZhY3Rpb249Y2xpY2tSZWRpcmVjdCZkb05vdExvZ0NsaWNrPXRydWU&th=1)
- [Circulation pump](https://www.amazon.com/Uniclife-80-550GPH-Submersible-Fountain-Hydroponic/dp/B00ZW6OHHY/ref=asc_df_B00ZW6OHHY/?tag=hyprod-20&linkCode=df0&hvadid=167149786275&hvpos=&hvnetw=g&hvrand=12117717222336054401&hvpone=&hvptwo=&hvqmt=&hvdev=c&hvdvcmdl=&hvlocint=&hvlocphy=9028280&hvtargid=pla-313113201776&psc=1)(s)

Tools List:

- Wire Strippers
- Skinny Body Philips head screwdriver
- Hot Glue Gun or Waterproof silicone calking

Software:

- [Arduino Software (IDE)](https://www.arduino.cc/en/software)
- [CoolTerm Data Recording](http://freeware.the-meiers.org/)
